# Cell type-specific intracellular protein delivery with inactivated botulinum toxin

**DOI:** 10.1101/2023.03.03.531050

**Authors:** Heegwang Roh, Brigitte G. Dorner, Alice Y. Ting

## Abstract

The ability to deliver proteins and peptides across the plasma membrane into the cytosol of living mammalian cells would be highly impactful for both basic science and medicine. Natural cell-penetrating protein toxins have shown promise as protein delivery platforms but existing approaches are limited by immunogenicity, lack of cell type specificity, or their multicomponent nature. Here we explore inactivated botulinum toxin (BoNT) as a protein delivery platform. Using split luciferase reconstitution in the cytosol as a readout for endosomal escape and cytosolic delivery, we showed that BoNT chimeras with nanobodies replacing their natural receptor binding domain could be selectively targeted to cells expressing nanobody-matched surface markers. We used chimeric BoNTs to deliver a range of cargo from 1.3-55 kD, and demonstrated selective delivery of orthogonal cargoes to distinct cell populations within a heterogenous mixture. These explorations suggest that BoNT can be a versatile platform for targeted protein and peptide delivery into mammalian cells.

## Introduction

Delivery of large macromolecules across the plasma membrane into the cytosol of mammalian cells remains a formidable challenge for both medicine and basic science. Antibodies and other biologics have well-established clinical utility against cell surface and extracellular targets, but have generally not been extended to intracellular targets due to their inability to cross the plasma membrane. In basic science, the use of large non-genetically encoded biophysical probes such as fluorophore-conjugated proteins or nanoparticles for the study of intracellular signaling has also been stymied by the lack of efficient methods for delivering such probes into the cytosol of living cells.

Because of the central importance of this problem, many different strategies for cytosolic delivery have been explored, and recent years have seen an explosion in lipid nanoparticle^1,2^ and engineered virus^3^ delivery platforms in particular. These technologies appear promising especially for the delivery of oligonucleotide-type cargo, although they largely lack cell type-specificity. For delivery of peptides and small proteins, cell-penetrating peptides^4,5^ and supercharged proteins^6,7^ have gained traction, although much of the cargo remains stuck in endosomes, causing toxicity and contributing background in some cases^8,9^.

Apart from these strategies, engineered toxins have emerged as a promising platform for the delivery of protein and peptide cargo into cells. A diverse array of toxins has evolved efficient strategies for receptor-mediated endocytosis followed by escape/translocation through endosomal membranes into the cytosol. Three major toxin classes have been explored as protein delivery platforms – anthrax toxin, diphtheria toxin, and botulinum neurotoxin – each with respective benefits and disadvantages. For example, the anthrax platform is modular and has been used to deliver Ras/Rap1-specific endopeptidase (RRSP) and the A-chain of diphtheria toxin using scFv-or IgG-mediated cell surface binding^10,11^. However, the multi-component nature of anthrax platform limits its utility and robustness. Engineered diphtheria toxin has been used to deliver a range of cargo (alpha-amylase^12^, purine nucleoside phosphorylase^13^, and RRSP^14^), but immunogenicity is a concern, as many individuals are vaccinated against diphtheria, and general reprogramming of cell type-specificity with artificial receptor binding domains has not yet been demonstrated.

We were intrigued by the botulinum neurotoxin (BoNT) platform in particular^15,16,17,18^, because people are not vaccinated against this toxin, reducing its immunogenicity; it is a single component system; and the toxin’s natural cargo – a protease that cleaves the synaptic vesicle fusion proteins SNAP25 or VAMP2 – has been re-engineered to cut alternative substrates^19^. Furthermore, a recent study utilizing BoNT for trans-synaptic tracing in flies^20^ suggests that the receptor binding domain of BoNT may be replaceable, opening the door to cell type-specific delivery. In this work, we explore BoNT as a protein delivery platform, developing a sensitive bioluminescence assay to detect cargo delivery into the cytosol. We demonstrate delivery of several distinct protein cargoes, and replace the receptor binding domain with nanobodies to achieve cell type-specificity.

## Results

BoNTs have three domains (**Figure 1A**, top): a receptor binding domain that enables receptor-mediated endocytosis, a translocation domain for escape from endosomes, and a zinc metalloprotease which is BoNT’s natural cargo. BoNT’s protease cuts either SNAP25 or VAMP2, which are both essential for synaptic vesicle fusion, thereby inhibiting synaptic transmission in the brain. Four different serotypes of BoNT (A, C, D, and X) have been used in protein delivery research^15,16,17,18^. For our study, we selected BoNT/X^21^, because its protease-dead triple mutant (E228Q/R360A/Y363F) has the least in vivo toxicity compared to other BoNT serotypes^16,22^.

**Figure 1.**
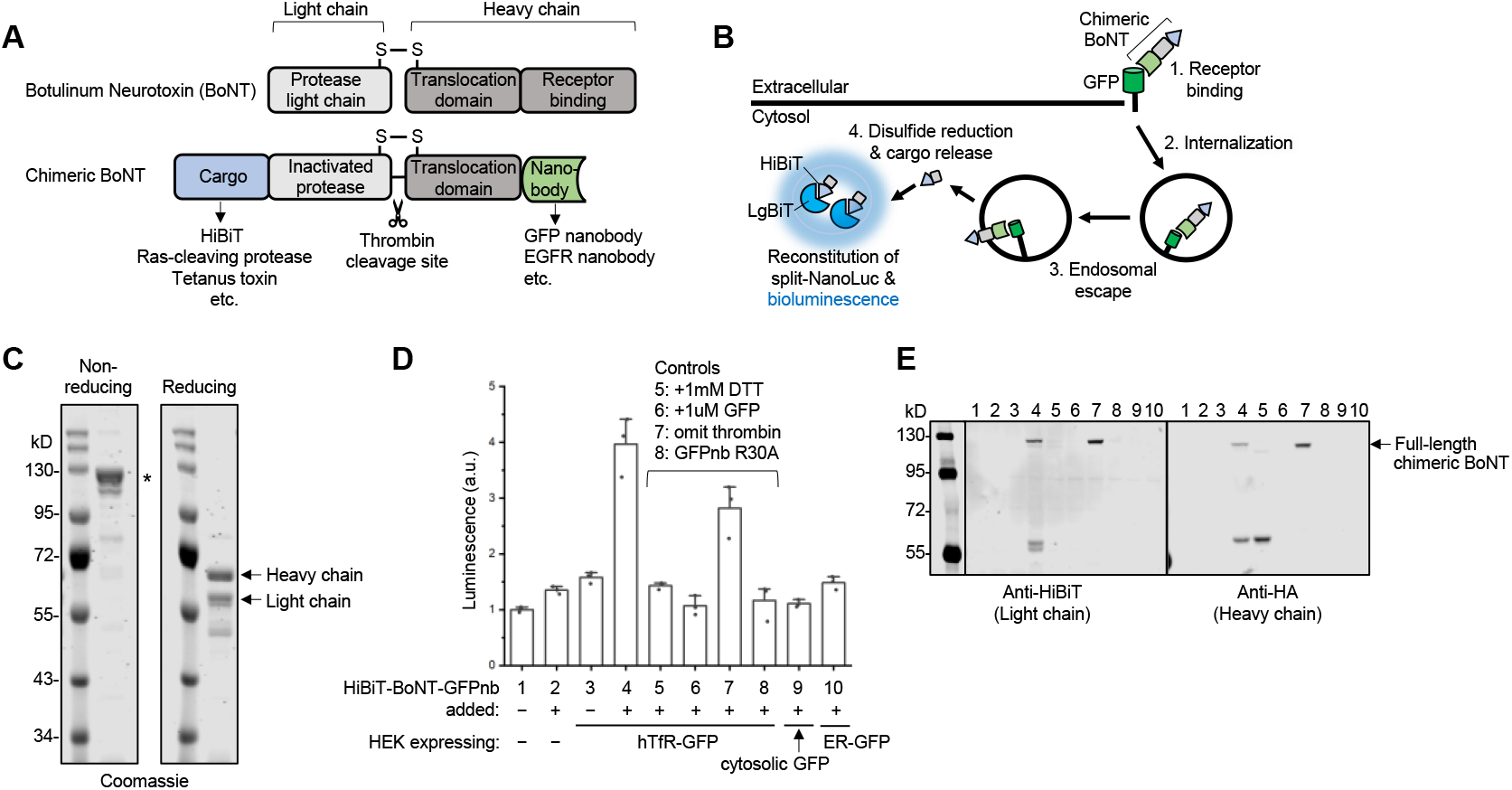
Design of chimeric botulinum toxins (BoNTs) and split-luciferase assay for detecting cytosolic cargo delivery. (**A**) Structure of wild-type BoNT and chimeric BoNTs. The light chain protease is inactivated by three mutations (E228Q/R360A/Y363F). The receptor binding domain is replaced by a nanobody. Protein cargo is fused at the N-terminus. (B)Schematic of chimeric BoNT binding to surface GFP-expressing mammalian cells, and delivering HiBiT cargo to the cytosol. HiBiT reconstitutes with cytosolically-expressed LgBiT to generate luciferase activity and bioluminescence in the presence of its substrate, furimazine. (C) Coomassie-stained SDS-PAGE gel of purified recombinant HiBiT-BoNT-GFPnb (GFP nanobody) with and without reduction by DTT. * denotes full-length HiBiT-BoNT-GFPnb. (**D**) Bioluminescence readout from living cells expressing the indicated constructs and treated with 10 nM of HiBiT-BoNT-GFPnb for 24 hours. All HEK293T cells expressed mCherry-LgBiT in the cytosol. (**E**) Western blot analysis of cell lysates from (D) under non-reducing conditions. Released HiBiT cargo (55 kD) is detected only in sample 4.

To begin, we sought to develop a highly specific and sensitive assay for BoNT-mediated delivery of cargo into the cytosol of living mammalian cells. Many previous assays require cell lysis^10,11,13,14,15,16,17^, which can produce artifactual results^23,24^ due to mixing of assay components post-lysis. We wished to develop a live-cell assay that could unambiguously distinguish between successful cytosolic delivery and trapping within endosomes. We also required high sensitivity to detect small quantities of delivered cargo. We selected the split-luciferase system NanoBiT for our assay^25^ because the fragments can be targeted to specific cellular subcompartments, reconstitution occurs rapidly and with high affinity (Kd 700 pM), and the bioluminescence readout is amplified and therefore highly sensitive.

To test the specificity of the NanoBiT reporter, we prepared HEK293T cells expressing the large fragment of NanoBiT, called LgBiT, in the cytosol. We also transduced the cells with the small fragment (HiBiT) targeted to the cytosol, endoplasmic reticulum (ER) lumen, or cell surface. As expected, bioluminescence was only detected for the cytosol co-localized combination, and not for cell surface-HiBiT or ER-HiBiT samples (**Figure S1**). This suggests that our assay will be able to faithfully detect cytosolic delivery of HiBiT fused to our cargo of interest (**Figure 1B**).

We then proceeded to design a BoNT variant in which the protease is inactivated, and 1.3 kD HiBiT is fused to its N-terminal end (**Figure 1A**, bottom). We also replaced the receptor binding domain of BoNT with a high-affinity (Kd 1 nM) nanobody against GFP (GFPnb)^26^. This engineered BoNT variant was expressed in *E. coli* and purified using a GST tag. The recombinant protein was treated with thrombin protease both to remove the GST tag and to cleave the toxin into “heavy chain” and “light chain” components connected via a disulfide bridge (**Figure 1A**, bottom). The SDS-PAGE gel in **Figure 1C** shows the full-length recombinant toxin (119 kD) and its reduced fragments (55 and 64 kD) after DTT treatment.

We prepared HEK293T cells stably expressing LgBiT-mCherry in the cytosol and introduced by transient transfection surface GFP fused to the transferrin receptor (hTfR-GFP), which is known to cycle constitutively through endosomes^20,27^. Incubation with 10 nM chimeric HiBiT-BoNT-GFPnb for 24 hours resulted in bioluminescence when furmazine, NanoBiT’s small-molecule substrate, was added to the live cells (**Figure 1D**). By contrast, bioluminescence was not detected when chimeric BoNT was not supplied, or cells did not express surface GFP. If the HEK293T cells expressed GFP in the cytosol or ER lumen instead of the cell surface, bioluminescence was also not detected (**Figure 1D**). To further confirm surface GFP-dependent entry, we either competed the toxin binding with an excess of recombinant GFP added to the extracellular medium, or we mutated the GFP nanobody portion of the toxin (R30A) (**Figure S2A**) to abolish its recognition of GFP^26^. Both treatments eliminated bioluminescence. Finally, we showed that reduction of the recombinant toxin with DTT to separate the cargo from the heavy chain prevented entry, but omission of the thrombin cleavage site (**Figure S2B**) did not, suggesting that a covalent link between heavy chain and cargo does not completely impair translocation and cytosolic entry^15, 21^. By comparing the bioluminescence of cells loaded with HiBiT-BoNT-GFPnb to the bioluminescence of purified recombinant NanoLuc, we estimate that our protocol results in delivery of at least 2.4 nM HiBiT cargo to the cytosol of live HEK293T cells (**Figure S3**; see **Supplementary Text** for calculations).

Western blot analysis (**Figure 1E**) of cellular samples from **Figure 1D** under non-reducing conditions showed that released HiBiT (55 kD) was detected only in sample 4, the surface GFP-expressing HEK293T cells treated with HiBiT-BoNT-GFPnb. Control samples did not uptake any toxin, with the exception of sample 7 which contained only full-length toxin due to omission of the thrombin cleavage site.

Collectively, these results show that NanoBiT can be used to read out BoNT-mediated cargo delivery to the cytosol, and that GFP:GFPnb recognition can mediate BoNT surface binding and entry into mammalian cells.

### Modularity of BoNT platform: Different cargos and different binders

To further explore the scope of BoNT-mediated protein delivery, we replaced the GFP nanobody in **Figure 1** with a 25 nM Kd nanobody against the extracellular domain of the epidermal growth factor receptor (EGFR)^28,29^ (EGFRnb; **Figure S2C**). EGFR is overexpressed in a number of human cancers (for example, >50% in triple-negative breast cancer^30^) and therefore a cell surface marker of great interest for targeted therapies^31,32^.

We carried out a dose-response experiment, measuring bioluminescence after incubating varying concentrations (0.1-100 nM) of recombinant toxin with HEK293T cells stably expressing cytosolic LgBiT and transiently expressing either surface GFP or EGFR (**Figures 2A-B**). Note that endogenous EGFR levels are low in HEK293T cells^33^. At 100 nM chimeric HiBiT-BoNT-EGFRnb, we detected ∼5-fold more uptake of HiBiT into EGFR-expressing cells than into GFP-expressing or untransfected cells, suggesting that the new EGFR nanobody mediates entry via binding to EGFR (**Figure 2B**). The reverse result was observed with our original HiBiT-BoNT-GFPnb toxin, which gave ∼4-fold more bioluminescence when incubated at 10 nM with surface GFP-expressing cells than with EGFR-expressing or untransfected cells (**Figure 2A**). The lower EC_50_ of the GFPnb toxin compared to the EGFRnb toxin (∼3 nM versus >10 nM) correlates with the reported binding affinities of their nanobody domains^26,28,29^.

**Figure 2.**
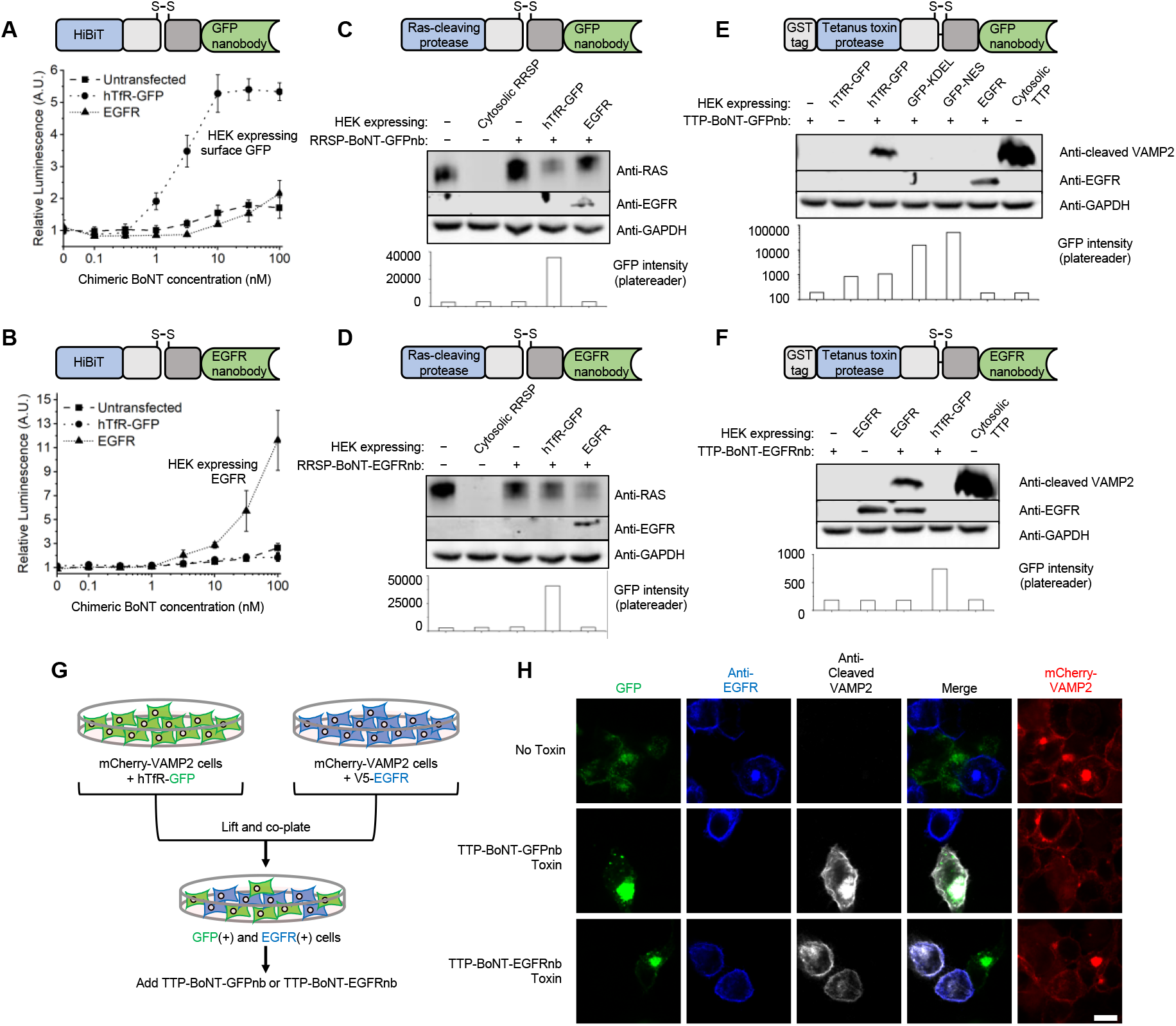
Chimeric BoNT delivers various cargo to mammalian cells in a receptor-dependent manner. (**A-B**) Delivery of HiBiT-BoNT-GFPnb (A) or HiBiT-BoNT-EGFRnb (B) chimeric toxins to HEK293T cells expressing surface GFP (hTfR-GFP) or EGFR. Bioluminescence reflects HiBiT (cargo) reconstitution with mCherry-LgBiT in the cytosol. (**C-D**) Delivery of Ras-cleaving protease (RRSP) to the cytosol of GFP-expressing or EGFR-expressing HEK293T cells, using chimeric toxins with GFPnb (C) or EGFRnb (D). HEK293T cells were incubated for 24 hours with 25 nM of RRSP-BoNT-GFPnb (C) or 100 nM of RRSP-BoNT-EGFRnb (D). Lysates were blotted with antibodies against pan-Ras, EGFR, and GAPDH (a cytosolic protein control). An expression plasmid encoding cytosolic RRSP was used as a positive control. GFP fluorescence was measured in live cells before cell lysis. (**E-F**) Delivery of tetanus toxin protease (TTP) to the cytosol of GFP-or EGFR-expressing cells, using chimeric toxins with GFPnb (E) or EGFRnb (F). All HEK293T cells expressed mCherry-VAMP2, whose cleavage was detected with anti-cleaved VAMP2 antibody. Cell were incubated for 24 hours with 100 nM of TTP-BoNT-GFPnb (E) or 100 nM of TTP-BoNT-EGFRnb (F). An expression plasmid encoding cytosolic TTP was used as a positive control. (**G**) Experimental design for chimeric BoNT delivery to a mixed culture of surface GFP-and EGFR-expressing VAMP2 reporter HEK293T cells. (**H**) Confocal fluorescence imaging of (G). Cells were treated with 100 nM of TTP-BoNT-GFPnb or TTP-BoNT-EGFRnb for 24 hours, and subsequently fixed and stained with anti-cleaved VAMP2 and anti-EGFR antibodies. In the second row, cleaved VAMP2 is only detected in GFP+ green cells, while in the third row, cleaved VAMP2 is only detected in EGFR+ blue cells. Scale bar, 10 μm.

Next, we explored alternative cargoes on the BoNT platform, beyond HiBiT. Previous studies using diphtheria toxin have delivered the Ras/Rap1-specific endopeptidase (RRSP), which cuts Ras and Rap1 between Y32 and D33 of the Switch I region, preventing downstream signaling^34,35^. RRSP is a therapeutically-relevant cargo, as constitutively active variants of Ras (HRAS, KRAS, and NRAS) are among the most common oncogenes^36,37^. We generated a chimeric toxin consisting of 55 kD RRSP fused to inactivated BoNT and GFP nanobody (RRSP-BoNT-GFPnb, **Figure S2D**). This material was incubated with HEK293T cells expressing either surface GFP or EGFR. After 24 hours, cells were lysed and blotted with anti-Ras antibody to quantify total remaining Ras protein. We observed the greatest Ras decrease (52%) in GFP-expressing HEK293T cells (**Figure 2C**), which matches the GFP nanobody of our engineered toxin. We also performed the reverse experiment, purifying toxin with EGFRnb in place of GFPnb (**Figure S2E**), and observed the greatest Ras decrease (72%) in EGFR-expressing rather than GFP-expressing HEK293T cells (**Figure 2D**).

For a third cargo, we utilized the protease of tetanus toxin (TTP), which cuts the synaptic vesicle fusion protein VAMP2. We fused 52 kD TTP to the N-terminal end of our chimeric BoNT, and GFPnb to the C-terminal end (**Figure S2F**). The recombinant toxin was incubated with HEK293T cells stably expressing mCherry-VAMP2 reporter, and 24 hours after treatment, the cells were lysed and blotted with an antibody that specifically detects cleaved VAMP2^38^. After treating cells with TTP-BoNT-GFPnb, cleaved VAMP2 was only detected in cells expressing surface GFP and treated with toxin, and not in control cells expressing cytosolic GFP or ER-localized GFP, nor in cells expressing EGFR (**Figure 2E**). We also prepared a TTP-BoNT-EGFRnb toxin (**Figure S2G**), which was selective for EGFR-expressing cells over surface GFP cells, as expected (**Figure 2F**).

To more rigorously assess the cell-type specificity of our TTP-BoNT chimeric toxins, we prepared a mixed culture of VAMP2 reporter cells expressing either surface GFP or EGFR (**Figure 2G**). When the mixed culture was treated with TTP-BoNT-GFPnb, only green GFP-positive cells showed staining with antibody against cleaved VAMP2, whereas neighboring GFP-negative cells did not (**Figure 2H** and **Figure S4**). Conversely, mixed cultures treated with TTP-BoNT-EGFRNb gave cleaved VAMP staining only on EGFR-positive cells. Thus, BoNT chimeras are able to deliver protein cargo to specific cell subpopulations within heterogeneous cultures.

### Simultaneous delivery of orthogonal cargoes to two different cell types in mixed culture

Encouraged by the cell type-specificity of the BoNT platform, we explored the possibility of delivering two different cargoes to two different cell types at the same time, by using orthogonal nanobodies that recognize distinct cell surface markers. We selected TTP as our first cargo, and the BoNT/A protease (BTP) as our second cargo. BTP from the BoNT/A serotype cleaves a different presynaptic protein^39^ (SNAP25) than the native protease of our BoNT/X-based platform, which cleaves VAMP2.

We prepared a mixed culture of HEK293T cells expressing either surface GFP or EGFR (**Figure 3A**). All cells also expressed both myc-VAMP2 reporter and FLAG-SNAP25 reporter (**Figure S5**). We then delivered a mixture of GFP-targeting TTP-BoNT-GFPnb toxin and EGFR-targeting BTP-BoNT-EGFRnb toxin. Cells were fixed and stained with antibodies against cleaved VAMP2 and cleaved SNAP25. GFP-positive cells showed cleaved VAMP2 staining, consistent with delivery of TTP cargo, while EGFR-positive cells showed cleaved SNAP25 staining, consistent with delivery of BTP cargo (**Figure 3B** and **Figure S6**). These results illustrate the high versatility and specificity of the BoNT platform for delivery of different protein cargos to selective cell populations.

**Figure 3.**
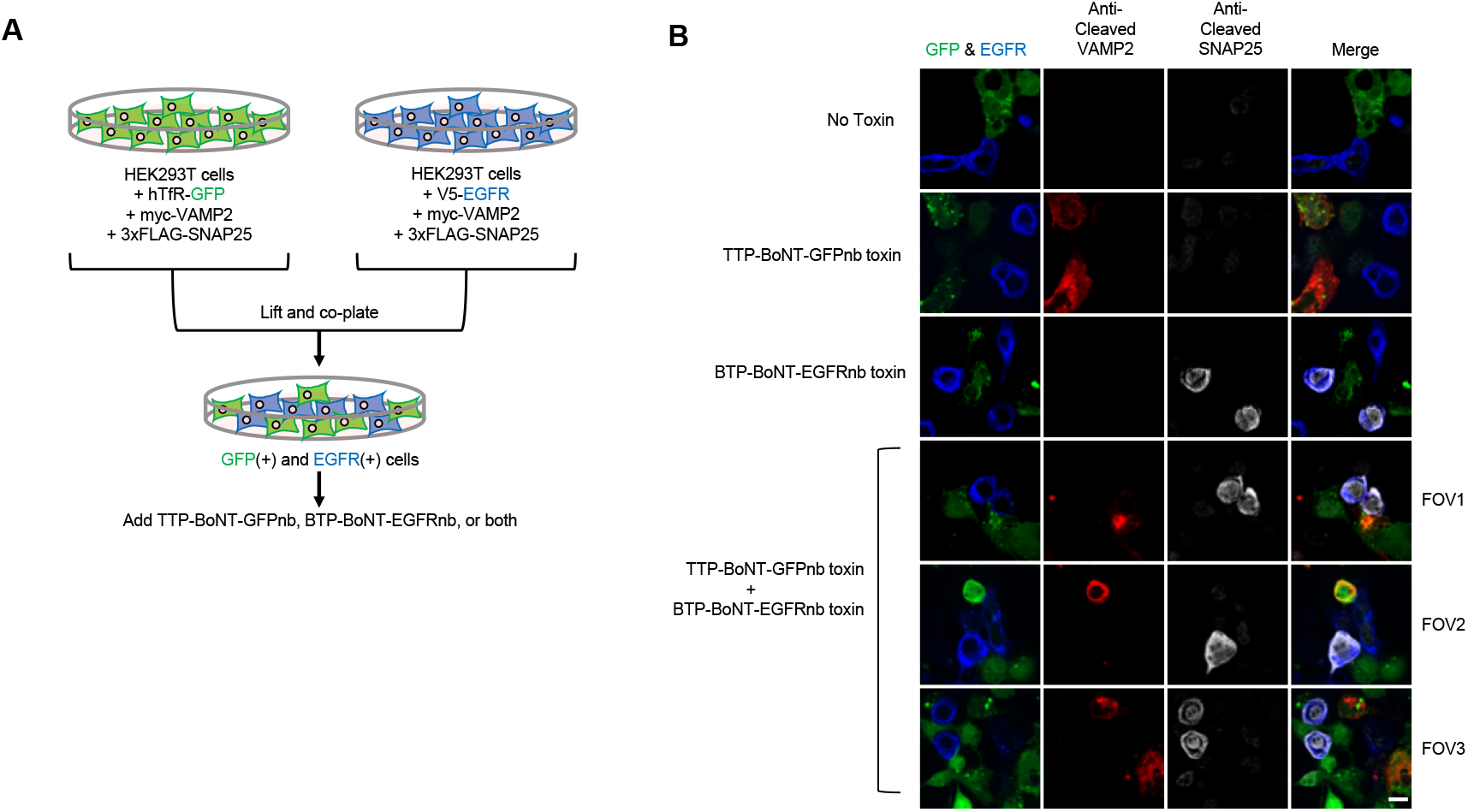
Simultaneous delivery of orthogonal toxins to two different cell types in an *in vitro* mixed culture system. (**A**) Experimental design. All HEK293T cells expressed both FLAG-SNAP25 and myc-VAMP2 reporters. (**B**) Confocal fluorescence imaging of (A). Cells were treated with 50 nM of either TTP-BoNT-GFPnb, BTP-BoNT-EGFRnb, or both for 24 hours, and subsequently fixed and stained with anti-cleaved VAMP2, anti-cleaved SNAP25, and anti-EGFR antibodies. In the bottom three rows, showing three different fields of view (FOV), only blue EGFR+ cells show anti-SNAP25 staining, evidence of BTP delivery. Only green GFP+ cells show anti-cleaved VAMP2 staining, evidence of TTP delivery. Scale bar, 10 μm.

## Discussion

In this work, we have shown that BoNT is a versatile platform for cytosolic delivery of protein cargoes in a cell-type specific manner. We showed that BoNT’s receptor binding domain can be replaced by two different nanobody binders, and we have demonstrated delivery of four different protein cargoes (HiBiT, RRSP, TTP, BTP) ranging in size from 1.3-55 kD. Our NanoBiT-based live cell assay provides a specific, sensitive, and somewhat quantitative readout of cytosolic cargo delivery, which may also have utility for other protein delivery platforms.

Our study used nanobodies for cell surface marker recognition, while other studies have used RBDs from other toxins^16,18^ or antibody-type binders that require chemical conjugation^10,11^. Nanobodies are straightforward to introduce to chimeric toxins by genetic fusion, and are available for a wide array of cell surface proteins^40^.

The sensitivity of the BoNT platform is comparable to but not superior to other toxins. Here we used 10-100 nM of chimeric toxin, which is within range of concentrations used in other BoNT studies (100 pM-3 nM^16^ or 250 nM-1 uM^18^ for example). Other toxin delivery platforms have used similar toxin concentrations as well (300 pM-100 nM for anthrax toxin^10,11^ and 10 pM-200 nM for diptheria toxin^12,13,14^).

Though BoNT has impressive capabilities, we have also discovered significant limitations to this delivery platform. Several recombinant BoNT fusions that we made in *E. coli* went to inclusion bodies, or precipitated after thrombin cleavage, suggesting low solubility. Cargo delivery yield is also limited; though we estimate that ∼2.4 nM of our smallest cargo, 1.3 kD HiBiT, could be delivered, larger cargoes are probably delivered in much smaller quantities. For this reason, enzyme cargoes such as RRSP, TTP, and BTP are attractive because small delivered quantities can exert detectable effects due to catalysis and signal amplification. We attempted but failed to deliver the large non-enzymatic cargoes split-GFP and Gal4 (data not shown).

Several lines of future work could improve the utility and robustness of the BoNT platform. First, a systematic study of the relationship between cargo size, charge, stability and the cytosolic delivery efficiency would aid users in cargo selection and design. Some studies have suggested that the ease with which a cargo can be unfolded and refolded is a major determinant of delivery efficiency^15,41^. Indeed, BoNT’s natural cargo, the 52 kD metalloprotease that cleaves VAMP2 or SNAP25, is metastable, with a low Tm of 48 °C^42^. In addition, protein engineering to improve BoNT chimera solubility, especially with the aid of computational algorithms such as PROSS^43^ could also improve the success rate of chimeric BoNTs.

## Author contributions

H.R. and A.Y.T. designed the research and analyzed all the data except where noted. H.R. performed all experiments. B.G.D. provided the anti-cleaved VAMP2 antibody. H.R. and A.Y.T. wrote the paper.

## Notes

The safety protocol for this study was reviewed by Stanford Administrative Panel on Biosafety (APB protocol #3902). All proteins were purified in quantities less than 0.1 mg.

H.R., B.G.D, and A.Y.T. declare no competing financial interest.

## Acknowledgements

This work was supported by the Chan Zuckerberg Biohub, the Stanford Wu Tsai Neurosciences Institute, and Stanford. A.Y.T. is a Chan Zuckerberg Biohub investigator.

