## Supplementary for "Cell type-specific intracellular protein delivery with inactivated botulinum toxin"

for

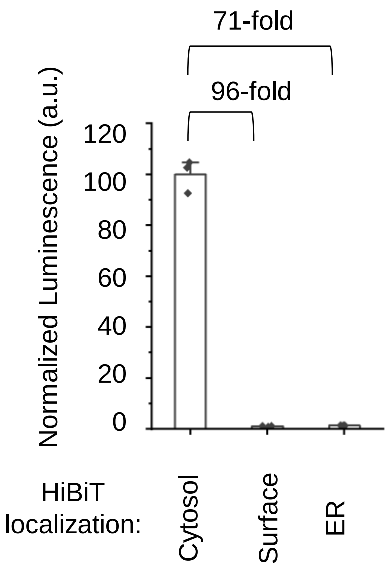


**Supplementary Figure 1. HiBiT and LgBiT must be co-compartmentalized to give bioluminescence.** HEK293T cells stably expressing cytosolic mCherry-LgBiT were transfected with plasmids encoding HiBiT targeted to the cytosol, cell surface, or ER lumen. Only cells expressing cytosolic HiBiT showed high bioluminescence upon addition of furimazine.

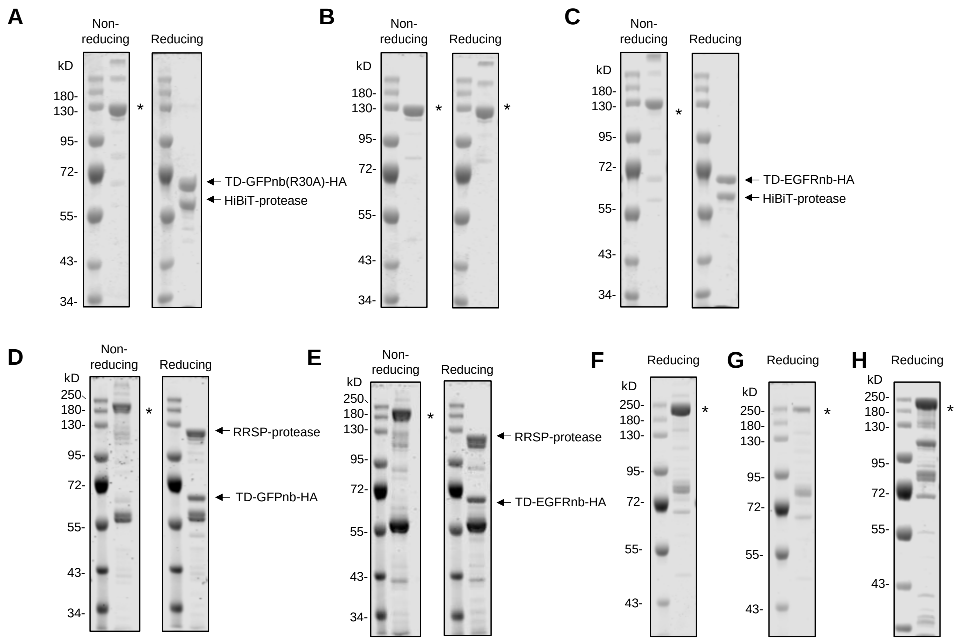


**Supplementary Figure 2. Additional data on purification and characterization of chimeric BoNTs.** Coomassie-stained SDS-PAGE gels of purified recombinant: (**A**) HiBiT-BoNT-GFPnb(R30A), (**B**) HiBiT-BoNT-GFPnb (single chain, no thrombin cleavage site), (**C**) HiBiT-BoNT-EGFRnb, (**D**) RRSP-BoNT-GFPnb, (**E**) RRSP-BoNT-EGFRnb, (**F**) TTP-BoNT-GFPnb, (**G**) TTP-BoNT-EGFRnb, (**H**) BTP-BoNT-EGFRnb. For chimeric BoNTs with TTP or BTP as cargo, thrombin cleavage was omitted and single-chain toxin was used due to the poor solubility of the di-chain form. *s denote full-length chimeric BoNTs. TD, translocation domain. HA, hemagglutinin epitope tag.

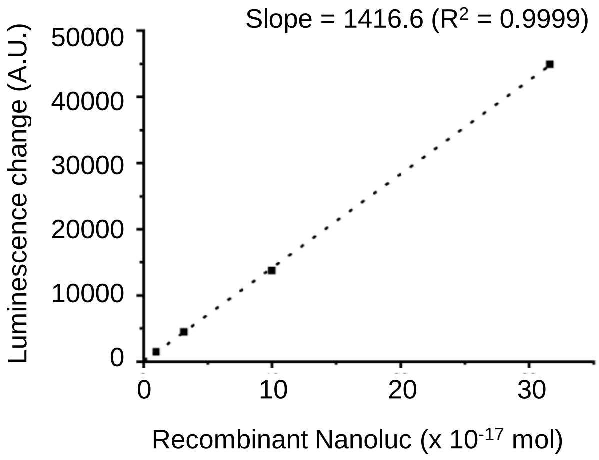


| | #1 | #2 | #3 |
|---|---|---|---|
| Luminescence change | 14498 | 18664 | 17891 |
| Nanoluc amount (mol) | 1.02 x $10^{-16}$ | 1.32 x $10^{-16}$ | 1.26 x $10^{-16}$ |
| Cell count | 4.32 x $10^{4}$ | 5.52 x $10^{4}$ | 5.11 x $10^{4}$ |
| Total cell volume (L) | 4.32 x $10^{-8}$ | 5.52 x $10^{-8}$ | 5.11 x $10^{-8}$ |
| Concentration (mol/L) | 2.37 x $10^{-9}$ | 2.39 x $10^{-9}$ | 2.47 x $10^{-9}$ |
| | **2.37 nM** | **2.39 nM** | **2.47 nM** |

**Supplementary Figure 3. Estimation of delivered cytosolic HiBiT concentration using recombinant Nanoluc standard curve.** Luminescence increase in mCherry-LgBiT HEK293T cells treated with 10 nM of of HiBiT-BoNT-GFPnb for 24 hours. Standard curve with purified recombinant Nanoluc shown above. See Methods for details.

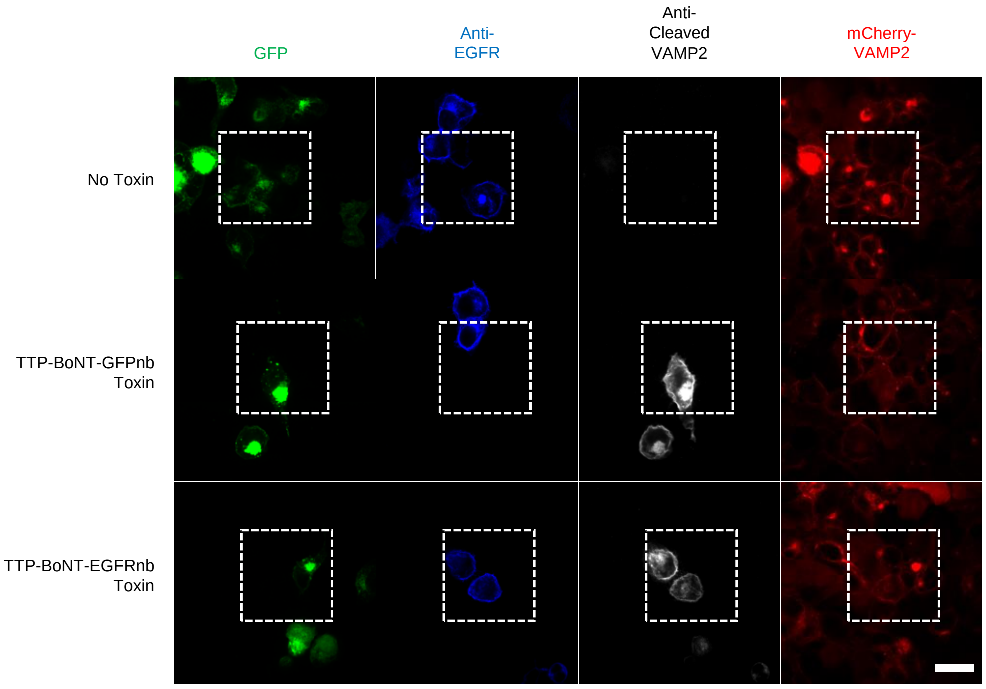


**Supplementary Figure 4. Additional data on chimeric BoNT-mediated delivery of TTP.** Related to Figure 2. Wider fields-of-view for images shown in Figure 2H. Scale bar, 20 μm.

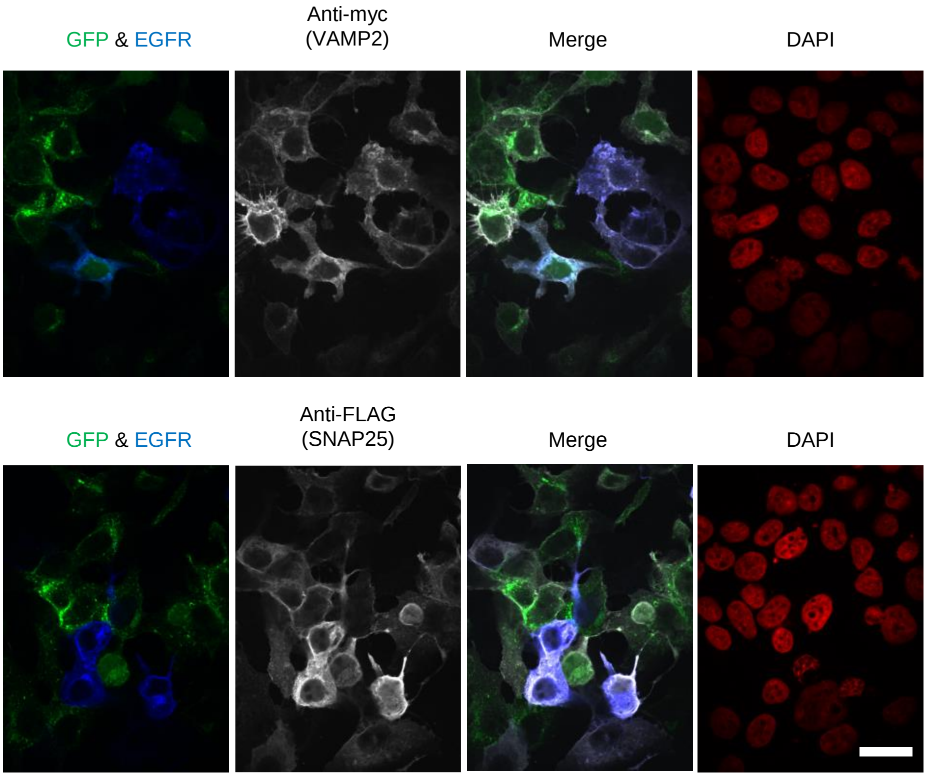


**Supplementary Figure 5. Both GFP-expressing and EGFR-expressing cells in the experiment in Figure 3 also express SNAP25 and VAMP2 reporters.** Related to Figure 3. The same batch of fixed cells from Figure 3 was stained with anti-myc or anti-FLAG antibody to detect myc-VAMP2 or FLAG-SNAP25, respectively. GFP-expressing and EGFR-expressing cells express both VAMP2 and SNAP25. Scale bar, 20 μm.

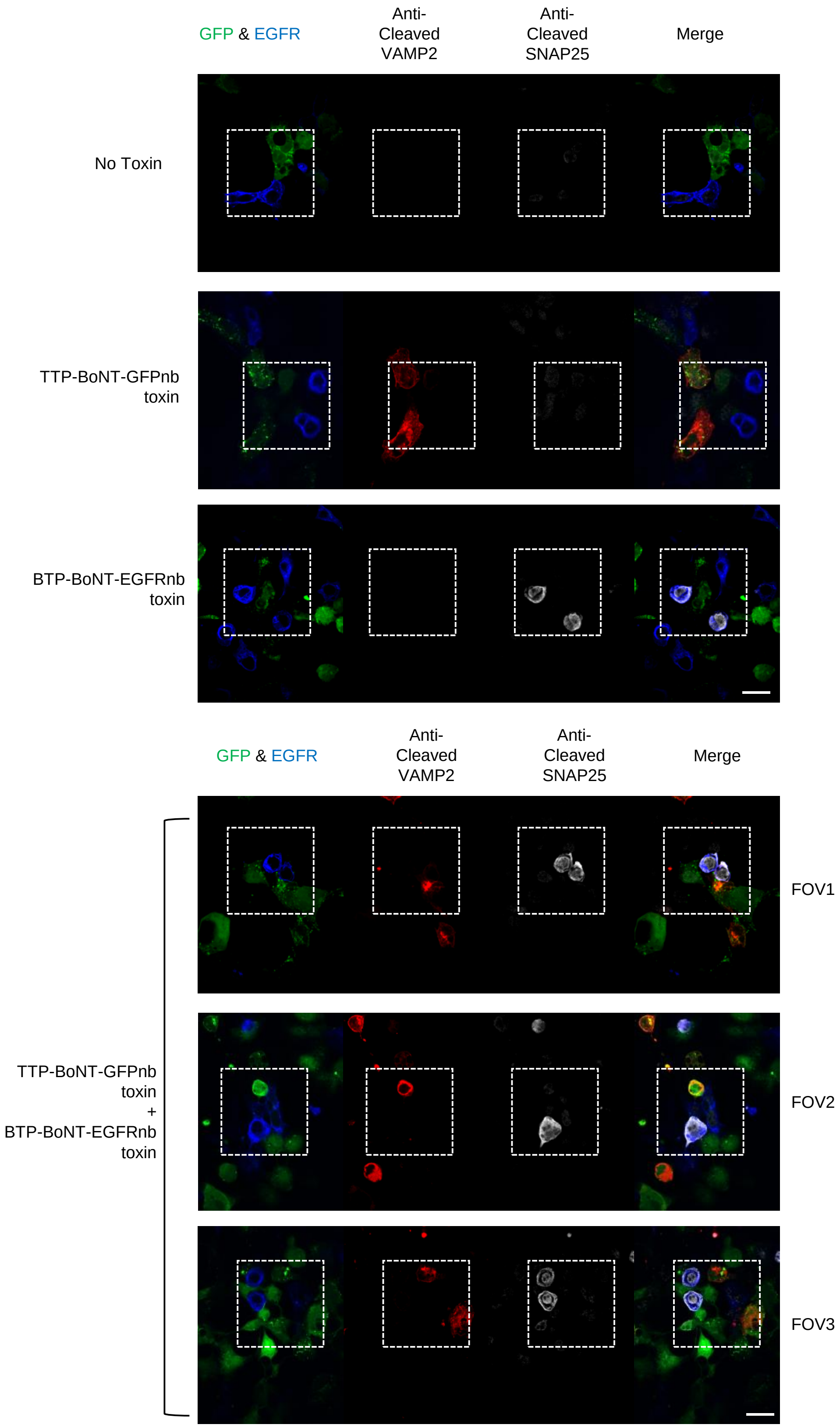


**Supplementary Figure 6. Additional data on the simultaneous delivery of TTP and BTP.** Related to Figure 3. Wider fields-of-view for images shown in Figure 3B. Scale bar, 20 μm.

### Materials

Table of plasmids used in this study.

| Plasmid name | Plasmid vector | Promoter | Features | Variants | Details | Used for | Used in |
|---|---|---|---|---|---|---|---|
| P1 | pLX304 | CMV | mCherry-16aa linker-LgBiT-18aa linker-FLAG-NES | | 16aa linker: SGSETPGTSESATPES; 18aa linker: GNGNGNGNGNGNGNGNGN; FLAG: DYKDDDDK; NES: LQLPPLERLTLD | Stable expression in HEK293T cells | **Figure 1, S1** |
| P2-3 | pDisplay | CMV | NheI-HiBiT-(GSSG)2-BoNT/X LC-Target motif-NotI | Target motif: CD4 TM, KDEL | BoNT/X contains a triple mutation (E228Q/R360A/Y363F) that inactivates the protease activity[1] | Transient expression | **Figure S1** |
| P4 | pcDNA | CMV | NheI-HiBiT-(GSSG)2-BoNT/X LC-6aa linker-NES-XhoI | | BoNT/X contains a triple mutation (E228Q/R360A/Y363F) that inactivates the protease activity[1]; 6aa linker: KSLVPR; NES: LQLPPLERLTLD | Transient expression | **Figure S1** |
| P5-6 | pGEX | Tac | BamHI-HiBiT-(GSSG)2-BoNT/X LC-Thrombin site-BoNT/X TD-Nanobody-HA-HindIII | Nanobody: GFPnb, EGFRnb | See section "Protein Sequences" | Overexpression in *E. coli* | **Figure 1, S2, S3** |
| P7 | pAAV | CMV | HindIII-hTfR-Syb-EGFP-EcoRI | | See reference[2] for details | Transient expression | **Figure 1, S3** |
| P8 | pcDNA | CMV | NotI-EGFP-NES-XhoI | | NES: LQLPPLERLTLD | Transient expression | **Figure 1** |
| P9 | pDisplay | CMV | IgK-7aa linker-EGFP-3aa linker-KDEL | | 7aa linker: GAQPARS; 3aa linker:GSG | Transient expression | **Figure 1** |
| P10 | pET21a | T7 | NdeI-EGFP-XhoI | | Includes C-terminal 6xHis tag | Overexpression in *E. coli* | **Figure 1** |
| P11 | pET | T7 | NcoI-6xHis-TEVcs-Nanoluc-BamHI | | Backbone: https://www.addgene.org/29659/; TEVcs: ENLYFQGS | Overexpression in *E. coli* | **Figure S3** |
| P12 | pAAV | CMV | HindIII-IL2ss-V5-EGFR-XhoI | | V5: GKPIPNPLLGLDST | Transient expression | **Figure 2** |
| P13-14 | pGEX | Tac | BamHI-RRSP-(GSSG)2-V5-(GSSG)2-BoNT/X LC-Thrombin site-BoNT/X TD-Nanobody-HA-HindIII | Nanobody: GFPnb, EGFRnb | See section "Protein Sequences" | Overexpression in *E. coli* | **Figure 2, S2** |
| P15-16 | pGEX | Tac | BamHI-V5-12aa linker-TTP-(GSSG)2-BoNT/X LC-Thrombin site-BoNT/X TD-Nanobody-HA-HindIII | Nanobody: GFPnb, EGFRnb | See section "Protein Sequences" | Overexpression in *E. coli* | **Figure 2, S2, S4, S5** |
| P17 | pLX208 | CMV | NheI-mCherry-6aa linker-myc-4aa linker-VAMP2-MluI | | 6aa linker: ASVNTG; 4aa linker: TGLK | Stable expression in HEK293T cells | **Figure 2, S4** |
| P18 | pGEX | Tac | BamHI-BTP-(GSSG)2-BoNT/X LC-Thrombin site-BoNT/X TD-EGFRnb-HA-HindIII | | See section "Protein Sequences" | Overexpression in *E. coli* | **Figure 3, S2, S5** |
| P19 | pAAV | CMV | 3xFLAG-SNAP25 | | 3xFLAG: DYKDHDGDYKDHDIDYKDDDDKH | Transient expression | **Figure 3, S5** |
| P20-22 | pcDNA | CMV | NotI-V5-12aa linker-Cargo-XhoI | Cargo: RRSP, TTP, BTP | 12aa linker: KGSGSTSGSGTG | Transient expression | **Figure 2, 3** |
| P23 | pLX208 | CMV | myc-4aa linker-VAMP2 | | 4aa linker: TGLK | Transient expression | **Figure 3, S5** |

Table of antibodies used in this study.

| Antibody | Source | Vendor | Catalog Number | Dilution(s) |
|---|---|---|---|---|
| Anti-HiBiT | Mouse | Promega | Clone 30E5 | WB: 1:1000 |
| Anti-V5 | Mouse | Thermo Fischer Scientific | R96025 | WB: 1:5000 |
| Anti-HA | Rabbit | Cell Signaling Technology | C29F4 | WB: 1:3000; IF: 1:1000 |
| Anti-myc | Mouse | Santa Cruz Biotechnology | Sc-40 | IF: 1:1000 |
| Anti-FLAG | Mouse | Abcam | Ab72469 | IF: 1:1000 |
| Anti-cleaved VAMP2 | Mouse | - | - | WB: 1:1000; IF: 1:1000 |
| Anti-GAPDH-HRP | Mouse | Santa Cruz Biotechnology | Sc-47724 HRP | WB: 1:3000 |
| Anti-RAS | Mouse | EMD Millipore | OP40 | WB: 1:1000 |
| Anti-EGFR | Rabbit | Cell Signaling Technology | 4267 | WB: 1:1000; IF: 1:1000 |
| Anti-cleaved SNAP25 | Mouse | GeneTex | B372M | WB: 1:1000; IF: 1:1000 |
| Anti-rabbit-AlexaFluor647 | Rabbit | Thermo Fischer Scientific | A21245 | IF: 1:1000 |
| Anti-mouse-AlexaFluor405 | Mouse | Thermo Fischer Scientific | A31553 | IF: 1:1000 |
| Anti-mouse-IRDye 800CW | Mouse | LI-COR Biosciences | 92632210 | IF: 1:3000 |
| Anti-rabbit-IRDye 680CW | Rabbit | LI-COR Biosciences | 92668071 | IF: 1:3000 |

### Methods

#### Cloning

For chimeric BoNT/X constructs, gene fragments were ordered from Twist Biosciences (see protein sequences under "Protein Sequences" below) and were subcloned into a pGEX-KG backbone (https://www.addgene.org/vector-database/2890/) using BamHI and HindIII restriction sites. All constructs were generated using standard cloning techniques. PCR fragments were amplified using Q5 polymerase (NEB Cat# M0491S). Vectors were digested using enzymatic restriction digest and ligated to gel purified PCR products using Gibson assembly. Ligated plasmid products were transformed into competent XL1-Blue *E. coli*.

#### Expression and purification of chimeric BoNTs from *E. coli*

Plasmids encoding chimeric BoNT/X constructs were used to transform *E. coli* BL21 (DE3) competent cells (Agilent Cat# 200131). Single colonies were used to inoculate 5 mL of Luria-Bertani broth (LB) supplemented with 100 mg/L ampicillin (LB-Amp) and grown overnight at 37 °C, 220 rpm. The overnight seed culture was diluted 1:100 into LB-Amp and was incubated at 37 °C, 220 rpm, until the $OD_{600}$ was between 0.3 and 0.6. The cells were then induced by adding IPTG to a final concentration of 0.2 mM and grown for approximately 18 hours at room temperature. After induction, cells were harvested by centrifugation at 8,000 g for 10 minutes.

For lysis, the cell pellet was frozen at -80 °C and thawed at room temperature to increase lysis efficiency. The cell pellet was solubilized by pipetting up and down in 2 mL lysis buffer per 100 mL of liquid culture. The lysis buffer is 20 mM HEPES 500 mM NaCl 5 mM EDTA 1 mM DTT supplemented with PMSF and 1X protease inhibitor cocktail (PIC, Thermo Scientific Cat# 78429). The lysate was pre-chilled on ice and sonicated using a Misonix sonicator (1 second on, 1 second off, for a total of 75 seconds on). The sonicated lysate was clarified by centrifugation for 10 minutes at 21,000 g, and the supernatant was filtered through a 0.45 µm filter.

Purification of GST-tagged chimeric toxins was performed using the Pierce™ GST spin purification kit (Pierce Cat# 16107), following manufacturer's protocols. Briefly, 4 mL of the clarified lysate was incubated with GST resin for at least 1 hour at 4 °C. After resin binding, the resin was washed four times with 4 mL of the wash/equilibration buffer (provided in the kit) by spinning for 2 minutes at 700 g. The bound protein was eluted twice using 1 mL of the elution buffer each time. Eluted proteins were exchanged into PBS using 30 kDa MWCO spin filter (Millipore Cat# UFC803024). Finally, eluted proteins were incubated with 10U/mg of Thrombin (Sigma-Aldrich Cat# T6634) overnight at 4 °C. After cleavage, residual thrombin and the cleaved GST tag in the mixture were removed using 50 kDa MWCO spin filter (Millipore Cat# UFC905024). It was important to elute the protein before carrying out thrombin cleavage, as on-resin thrombin cleavage resulted in a very low protein yield. For chimeric BoNT/X with TTP or BTP as the cargo, thrombin cleavage was omitted and single-chain toxin was used due to the poor solubility of the di-chain form. The resulting chimeric BoNT/X constructs were analyzed by SDS-PAGE under both reducing (10 mM DTT, boiled at 95 °C for 10 minutes) give time and temp if relevant) and non-reducing conditions, and stored in aliquots at -80 °C. Typical chimeric toxin protein yields were 1~5 mg/L.

Purification of 6xHis-tagged recombinant proteins was performed using Ni-NTA agarose beads. Briefly, the filtered lysate was batch loaded to 0.5 mL of packed Ni-NTA resin. The column was washed two times with 5 mL of wash buffer (50 mM Tris-HCl pH 8.0, 300 mM NaCl, 30 mM imidazole). The protein was then eluted using a total of 4 mL of elution buffer (50 mM Tris-HCl pH 8.0, 300 mM NaCl, 200 mM imidazole). The fractions were analyzed using SDS-PAGE, and the fractions containing the correct molecular weight were collected.

#### Cell culture and transfection

HEK293T cells from ATCC (<30 passages) were cultured as a monolayer in complete media (Dulbecco's modified Eagle's medium (DMEM, Gibco Cat# 11965-092) supplemented with 10% (w/v) fetal bovine serum (FBS, VWR Cat# 97068-085) and 1% penicillin-streptomycin (VWR Cat# 16777-164)) at 37 °C under 5% $CO_2$. In all experiments, glass coverslips and plates were pre-treated with 10 mg/mL human fibronectin (Millipore Cat# FC010) in Dulbecco's PBS (DPBS, Gibco Cat# 14190-144) for 30 minutes at room temperature before cell plating. For transient expression, cells were transfected at 50-70% confluency with indicated expression plasmids using PEI (Polysciences Cat# 24765-1) in DMEM without FBS. For cell cultures in 96-, 48-, 24-, and 6-well plates, 33, 100, 200, and 1000 ng of expression plasmids, respectively, were used for transfection.

#### Generation of mCherry-LgBiT and mCherry-VAMP2 reporter cell lines

Reporter cell lines were generated by lentiviral transduction and antibiotic selection. For lentivirus generation, HEK293T cells were cultured in 6-well plates and were transfected at approximately 70% confluency with 1000 ng of the lentiviral vector of interest, packaging plasmids pCMV-dR8.91 (900 ng), and pCMV-VSV-G (100 ng) with 12 µL of PEI. Approximately 48 hours after transfection, the cell medium was collected, filtered through a 0.45 µm filter, and then stored in 200 µL aliquots ("supernatant lentivirus"). The aliquots were flash-frozen in liquid nitrogen and stored at -80 °C.

For lentiviral transduction, HEK293T cells (<10 passages) were cultured in 6-well plates and transduced with 200 µL of supernatant lentivirus at approximately 50% confluency. Two days after transduction, cells with stably integrated reporters were selected by maintaining the cells in complete media containing antibiotics (8 µg/mL blasticidin for mCherry-LgBiT and 250 µg/mL hygromycin for mCherry-myc-VAMP2). Antibiotic selection was performed until all cells showed mCherry signal under a tabletop fluorescence microscope. After selection, reporter cell lines were grown in T75 tissue culture flask until confluency, and frozen stocks were prepared and stored in liquid nitrogen for future use.

##### Live-cell HiBiT:LgBiT complementation assay

For live-cell HiBiT:LgBiT complementation experiments, mCherry-LgBiT reporter cells were grown in 96-well plates and transfected with expression plasmids. 24 hours after transfection, the media was changed with 80 µL of complete media containing HiBiT toxin (0.1 to 100 nM). 24 hours after toxin treatment, cells were gently washed 3 times with complete media, and GFP fluorescence intensity was measured using a plate reader (TECAN M1000). Live-cell bioluminescence was then measured using Nano-Glo® Live Cell Assay System (Promega Cat# N2012), following manufacturer's protocols. Briefly, 1 volume of Nano-Glo® Live Cell Substrate with 19 volumes of Nano-Glo® LCS Dilution Buffer. This was further diluted 5-fold with complete media ("substrate mix"), and 80 µL of substrate mix was used each well.

##### Estimation of delivered cytosolic HiBiT concentration.

In this calculation, we make the assumption that reconstituted HiBiT:LgBiT complex has same luminescence intensity as full-length Nanoluc.

1. Measurement of luminescence from recombinant Nanoluc gave a standard curve of:
   Luminescence change = 1416.6 x ($10^{-17}$mol Recombinant Nanoluc)
2. For well #1, the luminescence increase compared to untreated cells was 14498, which means:
   (Amount HiBiT:LgBiT) = (14498) / (1416.6) x $10^{-17}$ mol = 1.02 x $10^{-16}$ mol
3. After resuspending the cells in DPBS, cell number was counted by tryphan blue staining:
   (Cell count) = 4.32 x $10^{4}$ cells
4. Assuming that one cell has a volume of 1 pL[3], the concentration of cytosolic HiBiT can be calculated as:
   (Amount HiBiT) / (Total cell volume) = (1.02 x $10^{-16}$ mol) / (4.32 x $10^{4}$ x $10^{-12}$ L) = 2.37 x $10^{-9}$ mol/L

##### Western blotting

In all western blot experiments, cells were transfected with expression plasmids and treated with toxin as described above (see "Live-cell HiBiT:LgBiT complementation assay"). 24 hours after toxin treatment, cells were gently washed three times with DPBS and lysed using 1X RIPA lysis buffer (Millipore Cat# 20-188) supplemented with PMSF and PIC. The concentrations of cell lysates were normalized using a Pierce™ BCA Protein Assay Kit (Pierce Cat# 23225) and samples were loaded onto a polyacrylamide gel (7 % gel for HiBiT toxin experiments, 12 % gel for RRSP and TTP toxin experiments).

After SDS-PAGE, the gels were transferred to a nitrocellulose membrane, and then stained with Ponceau S (0.1% (w/v) Ponceau S in 5 % acetic acid/water) for 1 minute to check protein loading. The Ponceau S stain was then washed out with 1X Tris-buffered saline with 0.1% Tween-20 (TBST, Teknova Cat# T1688). The blots were blocked in 5% (w/v) nonfat dry milk (LabScientific Cat# M8041) in TBST for at least 30 minutes at room temperature. The blots were washed three times with TBST for at least 5 minutes each time and then stained with primary antibodies in TBST overnight at 4 °C. The blots were washed three times with TBST and then stained with secondary antibodies (Anti-mouse-IRDye 800CW and Anti-rabibt-IRDye

680CW) in TBST for 1 hour at room temperature. The blots were washed three times with TBST and imaged using on a gel imager (Odyssey® CLx, LI-COR Biosciences).

#### Confocal fluorescence microscopy

For confocal fluorescence microscopy experiments, cells were grown on 12mm-diameter glass coverslips (VWR Cat# 72230-01) in 24-well plates. Cells were transfected with expression plasmids and treated with toxin as described above. 24 hours after toxin treatment, cells were gently washed three times with DPBS and were fixed with 4% (v/v) paraformaldehyde (PFA, ChemCruz Cat# sc-281692) at room temperature for 15 minutes. After fixation, PFA was aspirated and cells were permeabilized for 5 minutes with ice-cold methanol. Cells were washed again three times with DPBS and blocked for 1 hour with 1% bovine serum albumin (BSA, Fischer Scientific Cat# BP1600-1) in TBST at 4 °C. Cells were then incubated with primary antibodies in TBST overnight at 4 °C. After washing three times with TBST, cells were incubated with secondary antibodies (anti-mouse Alexa Fluor 647 and anti-rabbit Alexa Fluor 405) in TBST for 1 hour at room temperature. Cells were washed three times with TBST and imaged.

Imaging was performed with a Zeiss Axio Observer.Z1 microscope with a Yokogawa spinning disk confocal head, Cascade IIL:512 camera, a Quad-band notch dichroic mirror (405/488/568/647 nm), and 405 nm, 491 nm, 561 nm, and 640 nm lasers (all 50 mW). Images were captured through a 63× oil-immersion objective for the following fluorophores: AlexaFluor 405 (405 laser excitation, 445/40 emission), EGFP (491 laser excitation, 528/38 emission), mCherry and AlexaFluor 568 (561 laser excitation, 617/73 emission), and AlexaFluor 647 (647 laser excitation, 700/75 emission). Image acquisition times ranged from 10 to 500 ms per channel, and images were captured as the average of two or three such exposures in rapid succession. Image acquisition and processing was carried out with the SlideBook 5.0 software (Intelligent Imaging Innovations, 3i).

#### Delivery of RRSP and Ras cleavage assay

For delivery of RRSP using chimeric BoNT, HEK293T cells were grown in 24-well plates and transfected with expression plasmids. 24 hours after transfection, the media was changed with 350 µL of complete media containing RRSP toxin (25 to 100 nM). 24 hours after toxin treatment, cells were gently washed 3 times with DPBS and lysed. The cell lysate was used for Western blotting, as described above. Mouse Anti-Ras antibody and Rabbit Anti-EGFR antibody were used as primary antibodies, with 1:1000 dilution in TBST. After overnight staining, the blot was subsequently stained with Anti-mouse-IRDye 800CW and Anti-rabibt-IRDye 680CW, and imaged using a gel imager. After imaging, the blot was stained for 1 hour with HRP-conjugated anti-GAPDH antibody for 1 hour at room temperature. The blot was washed three times with TBST. The chemiluminescence was developed with Clarity™ Western ECL Substrate (Bio-Rad) and imaged using a ChemiDoc XRS (Bio-Rad) imaging system.

#### Delivery of TTP and VAMP2 cleavage assay

For delivery of TTP using chimeric BoNT, VAMP2 stable cells were grown in 24-well plates and transfected with expression plasmids. 24 hours after transfection, the media was changed with 350 µL of complete media containing 100 nM of TTP toxin. The subsequent cell lysis and Western blotting were carried out as previously described. For primary antibody staining, mouse anti-cleaved VAMP2 antibody was used with 1:1000 dilution.

The delivery of TTP was also assessed by confocal immunofluorescence imaging. HEK293T cells stably expressing mCherry-VAMP2 were cultured in 6-well plates and transfected with GFP-hTfR or EGFR expression plasmids. 12 hours after transfection, cells were lifted with Trypsin (Corning Cat# 25-053-CI) and a 1:1 mixture of the two cell populations was plated into new wells. 12 hours after co-plating, cells were treated with 100 nM of TTP toxins for 24 hours at 37 °C, and analyzed by confocal fluorescence microscopy as described above. Mouse anti-cleaved VAMP2 and Rabbit anti-EGFR antibodies were used as primary antibodies, and anti-rabbit-AlexaFluor 405 and anti-mouse-AlexaFluor647 antibodies were used as secondary antibodies.

**Simultaneous delivery of BTP and TTP**

For the simultaneous delivery of orthogonal BTP and TTP toxins, HEK293T cells were cultured in 6-well plates and transfected with GFP-hTfR or EGFR expression plasmids, as well as myc-VAMP2 and FLAG-SNAP25 expression plasmids. 12 hours after transfection, cells were lifted with Trypsin (Corning Cat# 25-053-CI) and a 1:1 mixture of the two cell populations was plated into new wells. 12 hours after co-plating, cells were treated with completed media containing 50 nM of TTP-BoNT-GFPnb and 50 nM of BTP-BoNT-EGFRnb, for 24 hours at 37 °C. Cells were fixed, stained, and confocal fluorescence microscopy as described above.

In this experiment, anti-cleaved SNAP25 and anti-cleaved VAMP2 antibodies were covalently conjugated with AlexaFluor 568 and AlexaFluor 647, respectively, as both antibodies are of mouse origin. Briefly, AlexaFluor568-NHS (Lumiprobe Cat#14820) and AlexaFluor647-NHS (Lumiprobe Cat#16820) were dissolved in DMSO. Antibody and AF-NHS ester were mixed in a 1:5 ratio and incubated for 2 hours at 4 °C. The reaction was quenched by diluting the mixture 20-fold in 20 mM Tris pH 8.0. Excess AF-NHS was removed by buffer exchanging the mixture into PBS using 30 kDa MWCO spin filter (Millipore Cat# UFC803024)

#### Protein sequences

**GST-HiBiT-BoNT/X-GFPnb**

MSPILGYWKIKGLVQPTRLLLEYLEEKYEEHLYERDEGDKWRNKKFELGLEFPNLPYYIDGDVKLTQS
MAIIRYIADKHNMLGGCPKERAEISMLEGAVLDIRYGVSRIAYSKDFETLKVDFLSKLPEMLKMFEDRL
CHKTYLNGDHVTHPDFMLYDALDVVLYMDPMCLDAFPKLVCFKKRIEAIPQIDKYLKSSKYIAWPLQG
WQATFGGGDHPPKSDLVPRGSVSGWRLFKKISGSSGGSSGMKLEINKFNYNDPIDGINVITMRPPRH
SDKINKGKGPFKAFQVIKNIWIVPERYNFTNNTNDLNIPSEPIMEADAIYNPNYLNTPSEKDEFLQGVIK
VLERIKSKPEGEKLLELISSSIPLPLVSNGALTLSDNETIAYQENNNIVSNLQANLVIYGPGPDIANNATY
GLYSTPISNGEGTLSEVSFSPFYLKPFDESYGNYRSLVNIVNKFVKREFAPDPASTLMHQ*LVHVTHNL
YGISNRNFYYNFDTGKIETSRQQNSLIFEELLTFGGIDSKAISSLIIKKIIETAKNNYTTLISERLNTVTVEN
DLLKYIKNKIPVQGRLGNFKLDTAEFEKKLNTILFVLNESNLAQRFSILVA*KHF*LKERPIDPIYVNILDDN
SYSTLEGFNISSQGSNDFQGQLLESSYFEKIESNALRAFIKICPRNGLLYNAIYRNSKNYLNNIDLEDKK
TTSKTNVSYPCSKSLVPRGSQALLNGCIEVENKDLFLISNKDSLNDINLSEEKIKPETTVFFKDKLPPQD
ITLSNYDFTEANSIPSISQQNILERNEELYEPIRNSLFEIKTIYVDKLTTFHFLEAQNIDESIDSSKIRVELT
DSVDEALSNPNKVYSPFKNMSNTINSIETGITSTYIFYQWLRSIVKDFSDETGKIDVIDKSSDTLAIVPYI
GPLLNIGNDIRHGDFVGAIELAGITALLEYVPEFTIPILVGLEVIGGELAREQVEAIVNNALDKRDQKWAE
VYNITKAQWWGTIHLQINTRLAHTYKALSRQANAIKMNMEFQLANYKGNIDDKAKIKNAISETEILLNKS
VEQAMKNTEKFMIKLSNSYLTKEMIPKVQDNLKNFDLETKKTLDKFIKEKEDILGTNLSSSLRRKVSIRL
NKNIAFDINDIPFSEFDDLINQYKNEIRSGMQVQLVESGGALVQPGGSLRLSCAASGFPVNRYSMRW
YRQAPGKEREWVAGMSSAGDRSSYEDSVKGRFTISRDDARNTVYLQMNSLKPEDTAVYYCNVNVG
FEYWGQGTQVTVSSKSGGYPYDVPDYA

Red: GST, Orange: Thrombin cleavage site, Blue: HiBiT, Gray: BoNT/X LC (*inactivating mutations), Dark Gray: BoNT/X TD, Green: GFPnb, Purple: HA

**GST-HiBiT-BoNT/X-GFPnb (single chain)**

MSPILGYWKIKGLVQPTRLLLEYLEEKYEEHLYERDEGDKWRNKKFELGLEFPNLPYYIDGDVKLTQS
MAIIRYIADKHNMLGGCPKERAEISMLEGAVLDIRYGVSRIAYSKDFETLKVDFLSKLPEMLKMFEDRL
CHKTYLNGDHVTHPDFMLYDALDVVLYMDPMCLDAFPKLVCFKKRIEAIPQIDKYLKSSKYIAWPLQG
WQATFGGGDHPPKSDLVPRGSVSGWRLFKKISGSSGGSSGMKLEINKFNYNDPIDGINVITMRPPRH
SDKINKGKGPFKAFQVIKNIWIVPERYNFTNNTNDLNIPSEPIMEADAIYNPNYLNTPSEKDEFLQGVIK
VLERIKSKPEGEKLLELISSSIPLPLVSNGALTLSDNETIAYQENNNIVSNLQANLVIYGPGPDIANNATY
GLYSTPISNGEGTLSEVSFSPFYLKPFDESYGNYRSLVNIVNKFVKREFAPDPASTLMHQ*LVHVTHNL
YGISNRNFYYNFDTGKIETSRQQNSLIFEELLTFGGIDSKAISSLIIKKIIETAKNNYTTLISERLNTVTVEN
DLLKYIKNKIPVQGRLGNFKLDTAEFEKKLNTILFVLNESNLAQRFSILVA*KHF*LKERPIDPIYVNILDDN
SYSTLEGFNISSQGSNDFQGQLLESSYFEKIESNALRAFIKICPRNGLLYNAIYRNSKNYLNNIDLEDKK
TTSKTNVSYPCSK**GSGSGS**QALLNGCIEVENKDLFLISNKDSLNDINLSEEKIKPETTVFFKDKLPPQ
DITLSNYDFTEANSIPSISQQNILERNEELYEPIRNSLFEIKTIYVDKLTTFHFLEAQNIDESIDSSKIRVEL
TDSVDEALSNPNKVYSPFKNMSNTINSIETGITSTYIFYQWLRSIVKDFSDETGKIDVIDKSSDTLAIVPYI
GPLLNIGNDIRHGDFVGAIELAGITALLEYVPEFTIPILVGLEVIGGELAREQVEAIVNNALDKRDQKWAE
VYNITKAQWWGTIHLQINTRLAHTYKALSRQANAIKMNMEFQLANYKGNIDDKAKIKNAISETEILLNKS
VEQAMKNTEKFMIKLSNSYLTKEMIPKVQDNLKNFDLETKKTLDKFIKEKEDILGTNLSSSLRRKVSIRL
NKNIAFDINDIPFSEFDDLINQYKNEIRSGMQVQLVESGGALVQPGGSLRLSCAASGFPVNRYSMRW
YRQAPGKEREWVAGMSSAGDRSSYEDSVKGRFTISRDDARNTVYLQMNSLKPEDTAVYYCNVNVG
FEYWGQGTQVTVSSKSGGYPYDVPDYA

Red: GST, Orange: Thrombin cleavage site, Blue: HiBiT, Gray: BoNT/X LC (*inactivating mutations), Dark Gray: BoNT/X TD, Green: GFPnb, Purple: HA

**GST-HiBiT-BoNT/X-GFPnb (R30A)**

MSPILGYWKIKGLVQPTRLLLEYLEEKYEEHLYERDEGDKWRNKKFELGLEFPNLPYYIDGDVKLTQS
MAIIRYIADKHNMLGGCPKERAEISMLEGAVLDIRYGVSRIAYSKDFETLKVDFLSKLPEMLKMFEDRL
CHKTYLNGDHVTHPDFMLYDALDVVLYMDPMCLDAFPKLVCFKKRIEAIPQIDKYLKSSKYIAWPLQG
WQATFGGGDHPPKSDLVPRGSVSGWRLFKKISGSSGGSSGMKLEINKFNYNDPIDGINVITMRPPRH
SDKINKGKGPFKAFQVIKNIWIVPERYNFTNNTNDLNIPSEPIMEADAIYNPNYLNTPSEKDEFLQGVIK
VLERIKSKPEGEKLLELISSSIPLPLVSNGALTLSDNETIAYQENNNIVSNLQANLVIYGPGPDIANNATY

GLYSTPISNGEGTLSEVSFSPFYLKPFDESYGNYRSLVNIVNKFVKREFAPDPASTLMHQ*LVHVTHNL
YGISNRNFYYNFDTGKIETSRQQNSLIFEELLTFGGIDSKAISSLIIKKIIETAKNNYTTLISERLNTVTVEN
DLLKYIKNKIPVQGRLGNFKLDTAEFEKKLNTILFVLNESNLAQRFSILVA*KHF*LKERPIDPIYVNILDDN
SYSTLEGFNISSQGSNDFQGQLLESSYFEKIESNALRAFIKICPRNGLLYNAIYRNSKNYLNNIDLEDKK
TTSKTNVSYPCSKSLVPRGSQALLNGCIEVENKDLFLISNKDSLNDINLSEEKIKPETTVFFKDKLPPQD
ITLSNYDFTEANSIPSISQQNILERNEELYEPIRNSLFEIKTIYVDKLTTFHFLEAQNIDESIDSSKIRVELT
DSVDEALSNPNKVYSPFKNMSNTINSIETGITSTYIFYQWLRSIVKDFSDETGKIDVIDKSSDTLAIVPYI
GPLLNIGNDIRHGDFVGAIELAGITALLEYVPEFTIPILVGLEVIGGELAREQVEAIVNNALDKRDQKWAE
VYNITKAQWWGTIHLQINTRLAHTYKALSRQANAIKMNMEFQLANYKGNIDDKAKIKNAISETEILLNKS
VEQAMKNTEKFMIKLSNSYLTKEMIPKVQDNLKNFDLETKKTLDKFIKEKEDILGTNLSSSLRRKVSIRL
NKNIAFDINDIPFSEFDDLINQYKNEIRSGMQVQLVESGGALVQPGGSLRLSCAASGFPVNRYSMA*W
YRQAPGKEREWVAGMSSAGDRSSYEDSVKGRFTISRDDARNTVYLQMNSLKPEDTAVYYCNVNVG
FEYWGQGTQVTVSSKSGGYPYDVPDYA

Red: GST, Orange: Thrombin cleavage site, Blue: HiBiT, Gray: BoNT/X LC (*inactivating mutations), Dark Gray: BoNT/X TD, Green: GFPnb (*R30A mutation), Purple: HA

**GST-HiBiT-BoNT/X-EGFRnb**

MSPILGYWKIKGLVQPTRLLLEYLEEKYEEHLYERDEGDKWRNKKFELGLEFPNLPYYIDGDVKLTQS
MAIIRYIADKHNMLGGCPKERAEISMLEGAVLDIRYGVSRIAYSKDFETLKVDFLSKLPEMLKMFEDRL
CHKTYLNGDHVTHPDFMLYDALDVVLYMDPMCLDAFPKLVCFKKRIEAIPQIDKYLKSSKYIAWPLQG
WQATFGGGDHPPKSDLVPRGSVSGWRLFKKISGSSGGSSGMKLEINKFNYNDPIDGINVITMRPPRH
SDKINKGKGPFKAFQVIKNIWIVPERYNFTNNTNDLNIPSEPIMEADAIYNPNYLNTPSEKDEFLQGVIK
VLERIKSKPEGEKLLELISSSIPLPLVSNGALTLSDNETIAYQENNNIVSNLQANLVIYGPGPDIANNATY
GLYSTPISNGEGTLSEVSFSPFYLKPFDESYGNYRSLVNIVNKFVKREFAPDPASTLMHQ*LVHVTHNL
YGISNRNFYYNFDTGKIETSRQQNSLIFEELLTFGGIDSKAISSLIIKKIIETAKNNYTTLISERLNTVTVEN
DLLKYIKNKIPVQGRLGNFKLDTAEFEKKLNTILFVLNESNLAQRFSILVA*KHF*LKERPIDPIYVNILDDN
SYSTLEGFNISSQGSNDFQGQLLESSYFEKIESNALRAFIKICPRNGLLYNAIYRNSKNYLNNIDLEDKK
TTSKTNVSYPCSKSLVPRGSQALLNGCIEVENKDLFLISNKDSLNDINLSEEKIKPETTVFFKDKLPPQD
ITLSNYDFTEANSIPSISQQNILERNEELYEPIRNSLFEIKTIYVDKLTTFHFLEAQNIDESIDSSKIRVELT
DSVDEALSNPNKVYSPFKNMSNTINSIETGITSTYIFYQWLRSIVKDFSDETGKIDVIDKSSDTLAIVPYI
GPLLNIGNDIRHGDFVGAIELAGITALLEYVPEFTIPILVGLEVIGGELAREQVEAIVNNALDKRDQKWAE
VYNITKAQWWGTIHLQINTRLAHTYKALSRQANAIKMNMEFQLANYKGNIDDKAKIKNAISETEILLNKS
VEQAMKNTEKFMIKLSNSYLTKEMIPKVQDNLKNFDLETKKTLDKFIKEKEDILGTNLSSSLRRKVSIRL
NKNIAFDINDIPFSEFDDLINQYKNEIQVKLEESGGGSVQTGGSLRLTCAASGRTSRSYGMGWFRQAP
GKEREFVSGISWRGDSTGYADSVKGRFTISRDNAKNTVDLQMNSLKPEDTAIYYCAAAAGSAWYGTL
YEYDYWGQGTQVTVSSSGGYPYDVPDYA

Red: GST, Orange: Thrombin cleavage site, Blue: HiBiT, Gray: BoNT/X LC (*inactivating mutations), Dark Gray: BoNT/X TD, Green: EGFRnb, Purple: HA

**GST-RRSP-V5-BoNT/X-GFPnb**

MSPILGYWKIKGLVQPTRLLLEYLEEKYEEHLYERDEGDKWRNKKFELGLEFPNLPYYIDGDVKLTQS
MAIIRYIADKHNMLGGCPKERAEISMLEGAVLDIRYGVSRIAYSKDFETLKVDFLSKLPEMLKMFEDRL
CHKTYLNGDHVTHPDFMLYDALDVVLYMDPMCLDAFPKLVCFKKRIEAIPQIDKYLKSSKYIAWPLQG
WQATFGGGDHPPKSDLVPRGSQELKERAKVFAKPIGASYQGILDQLDLVHQAKGRDQIAASFELNKK
INDYIAEHPTSGRNQALTQLKEQVTSALFIGKMQVAQAGIDAIAQTRPELAARIFMVAIEEANGKHVGLT
DMMVRWANEDPYLAPKHGYKGETPSDLGFDAKYHVDLGEHYADFKQWLETSQSNGLLSKATLDES
TKTVHLGYSYQELQDLTGAESVQMAFYFLKEAAKKADPISGDSAEMILLKKFADQSYLSQLDSDRMD
QIEGIYRSSHETDIDAWDRRYSGTGYDELTNKLASATGVDEQLAVLLDDRKGLLIGEVHGSDVNGLRF
VNEQMDALKKQGVTVIGLEHLRSDLAQPLIDRYLATGVMSSELSAMLKTKHLDVTLFENARANGMRIV
ALDANSSARPNVQGTEHGLMYRAGAANIAVEVLQNLPDGEKFVAIYGKAHLQSHKGIEGFVPGITH
RLDLPALKVSDSNQFTVEQDDVSLRVVYDDVANKPKITFKGSLGSSGGSSGGKPIPNPLLGLDSTGS
SGGSSGMKLEINKFNYNDPIDGINVITMRPPRHSDKINKGKGPFKAFQVIKNIWIVPERYNFTNNTNDL
NIPSEPIMEADAIYNPNYLNTPSEKDEFLQGVIKVLERIKSKPEGEKLLELISSSIPLPLVSNGALTLSDN
ETIAYQENNNIVSNLQANLVIYGPGPDIANNATYGLYSTPISNGEGTLSEVSFSPFYLKPFDESYGNYR
SLVNIVNKFVKREFAPDPASTLMHQ*LVHVTHNLYGISNRNFYYNFDTGKIETSRQQNSLIFEELLTFG

GIDSKAISSLIIKKIIETAKNNYTTLISERLNTVTVENDLLKYIKNKIPVQGRLGNFKLDTAEFEKKLNTILFV
LNESNLAQRFSILVA*KHF*LKERPIDPIYVNILDDNSYSTLEGFNISSQGSNDFQGQLLESSYFEKIESN
ALRAFIKICPRNGLLYNAIYRNSKNYLNNIDLEDKKTTSKTNVSYPCSKSLVPRGSQALLNGCIEVENKD
LFLISNKDSLNDINLSEEKIKPETTVFFKDKLPPQDITLSNYDFTEANSIPSISQQNILERNEELYEPIRNS
LFEIKTIYVDKLTTFHFLEAQNIDESIDSSKIRVELTDSVDEALSNPNKVYSPFKNMSNTINSIETGITSTYI
FYQWLRSIVKDFSDETGKIDVIDKSSDTLAIVPYIGPLLNIGNDIRHGDFVGAIELAGITALLEYVPEFTIPI
LVGLEVIGGELAREQVEAIVNNALDKRDQKWAEVYNITKAQWWGTIHLQINTRLAHTYKALSRQANAI
KMNMEFQLANYKGNIDDKAKIKNAISETEILLNKSVEQAMKNTEKFMIKLSNSYLTKEMIPKVQDNLKN
FDLETKKTLDKFIKEKEDILGTNLSSSLRRKVSIRLNKNIAFDINDIPFSEFDDLINQYKNEIRSGMQVQL
VESGGALVQPGGSLRLSCAASGFPVNRYSMRWYRQAPGKEREWVAGMSSAGDRSSYEDSVKGRF
TISRDDARNTVYLQMNSLKPEDTAVYYCNVNVGFEYWGQGTQVTVSSKSGGYPYDVPDYA

Red: GST, Orange: Thrombin cleavage site, Blue: RRSP, Gray: BoNT/X LC (*inactivating mutations), Dark Gray: BoNT/X TD, Green: GFPnb, Purple: HA, Brown: V5

**GST-RRSP-V5-BoNT/X-EGFRnb**

MSPILGYWKIKGLVQPTRLLLEYLEEKYEEHLYERDEGDKWRNKKFELGLEFPNLPYYIDGDVKLTQS
MAIIRYIADKHNMLGGCPKERAEISMLEGAVLDIRYGVSRIAYSKDFETLKVDFLSKLPEMLKMFEDRL
CHKTYLNGDHVTHPDFMLYDALDVVLYMDPMCLDAFPKLVCFKKRIEAIPQIDKYLKSSKYIAWPLQG
WQATFGGGDHPPKSDLVPRGSQELKERAKVFAKPIGASYQGILDQLDLVHQAKGRDQIAASFELNKK
INDYIAEHPTSGRNQALTQLKEQVTSALFIGKMQVAQAGIDAIAQTRPELAARIFMVAIEEANGKHVGLT
DMMVRWANEDPYLAPKHGYKGETPSDLGFDAKYHVDLGEHYADFKQWLETSQSNGLLSKATLDES
TKTVHLGYSYQELQDLTGAESVQMAFYFLKEAAKKADPISGDSAEMILLKKFADQSYLSQLDSDRMD
QIEGIYRSSHETDIDAWDRRYSGTGYDELTNKLASATGVDEQLAVLLDDRKGLLIGEVHGSDVNGLRF
VNEQMDALKKQGVTVIGLEHLRSDLAQPLIDRYLATGVMSSELSAMLKTKHLDVTLFENARANGMRIV
ALDANSSARPNVQGTEHGLMYRAGAANIAVEVLQNLPDGEKFVAIYGKAHLQSHKGIEGFVPGITH
RLDLPALKVSDSNQFTVEQDDVSLRVVYDDVANKPKITFKGSLGSSGGSSGGKPIPNPLLGLDSTGS
SGGSSGMKLEINKFNYNDPIDGINVITMRPPRHSDKINKGKGPFKAFQVIKNIWIVPERYNFTNNTNDL
NIPSEPIMEADAIYNPNYLNTPSEKDEFLQGVIKVLERIKSKPEGEKLLELISSSIPLPLVSNGALTLSDN
ETIAYQENNNIVSNLQANLVIYGPGPDIANNATYGLYSTPISNGEGTLSEVSFSPFYLKPFDESYGNYR
SLVNIVNKFVKREFAPDPASTLMHQ*LVHVTHNLYGISNRNFYYNFDTGKIETSRQQNSLIFEELLTFG
GIDSKAISSLIIKKIIETAKNNYTTLISERLNTVTVENDLLKYIKNKIPVQGRLGNFKLDTAEFEKKLNTILFV
LNESNLAQRFSILVA*KHF*LKERPIDPIYVNILDDNSYSTLEGFNISSQGSNDFQGQLLESSYFEKIESN
ALRAFIKICPRNGLLYNAIYRNSKNYLNNIDLEDKKTTSKTNVSYPCSKSLVPRGSQALLNGCIEVENKD
LFLISNKDSLNDINLSEEKIKPETTVFFKDKLPPQDITLSNYDFTEANSIPSISQQNILERNEELYEPIRNS
LFEIKTIYVDKLTTFHFLEAQNIDESIDSSKIRVELTDSVDEALSNPNKVYSPFKNMSNTINSIETGITSTYI
FYQWLRSIVKDFSDETGKIDVIDKSSDTLAIVPYIGPLLNIGNDIRHGDFVGAIELAGITALLEYVPEFTIPI
LVGLEVIGGELAREQVEAIVNNALDKRDQKWAEVYNITKAQWWGTIHLQINTRLAHTYKALSRQANAI
KMNMEFQLANYKGNIDDKAKIKNAISETEILLNKSVEQAMKNTEKFMIKLSNSYLTKEMIPKVQDNLKN
FDLETKKTLDKFIKEKEDILGTNLSSSLRRKVSIRLNKNIAFDINDIPFSEFDDLINQYKNEIQVKLEESGG
GSVQTGGSLRLTCAASGRTSRSYGMGWFRQAPGKEREFVSGISWRGDSTGYADSVKGRFTISRDN
AKNTVDLQMNSLKPEDTAIYYCAAAAGSAWYGTLYEYDYWGQGTQVTVSSSGGYPYDVPDYA

Red: GST, Orange: Thrombin cleavage site, Blue: RRSP, Gray: BoNT/X LC (*inactivating mutations), Dark Gray: BoNT/X TD, Green: EGFRnb, Purple: HA, Brown: V5

**GST-V5-TTP- BoNT/X-GFPnb**

MSPILGYWKIKGLVQPTRLLLEYLEEKYEEHLYERDEGDKWRNKKFELGLEFPNLPYYIDGDVKLTQS
MAIIRYIADKHNMLGGCPKERAEISMLEGAVLDIRYGVSRIAYSKDFETLKVDFLSKLPEMLKMFEDRL
CHKTYLNGDHVTHPDFMLYDALDVVLYMDPMCLDAFPKLVCFKKRIEAIPQIDKYLKSSKYIAWPLQG
WQATFGGGDHPPKSDLVPRGSGKPIPNPLLGLDSTKGSGSTSGSGTGPITINNFRYSDPVNNDTIIM
MEPPYCKGLDIYYKAFKITDRIWIVPERYEFGTKPEDFNPPSSLIEGASEYYDPNYLRTDSDKDRFLQT
MVKLFNRIKNNVAGEALLDKIINAIPYLGNSYSLLDKFDTNSNSVSFNLLEQDPSGATTKSAMLTNLIIFG
PGPVLNKNEVRGIVLRVDNKNYFPCRDGFGSIMQMAFCPEYVPTFDNVIENITSLTIGKSKYFQDPALL
LMHELIHVLHGLYGMQVSSHEIIPSKQEIYMQHTYPISAEELFTFGGQDANLISIDIKNDLYEKTLNDYKA
IANKLSQVTSCNDPNIDIDSYKQIYQQKYQFDKDSNGQYIVNEDKFQILYNSIMYGFTEIELGKKFNIKT
RLSYFSMNHDPVKIPNLLDDTIYNDTEGFNIESKDLKSEYKGQNMRVNTNAFRNVDGSGLVSKLIGLC

KKIIPPTNIRENLYNRTAGSSGGSSGMKLEINKFNYNDPIDGINVITMRPPRHSDKINKGKGPFKAFQVI
KNIWIVPERYNFTNNTNDLNIPSEPIMEADAIYNPNYLNTPSEKDEFLQGVIKVLERIKSKPEGEKLLELI
SSSIPLPLVSNGALTLSDNETIAYQENNNIVSNLQANLVIYGPGPDIANNATYGLYSTPISNGEGTLSEV
SFSPFYLKPFDESYGNYRSLVNIVNKFVKREFAPDPASTLMHQ*LVHVTHNLYGISNRNFYYNFDTGKI
ETSRQQNSLIFEELLTFGGIDSKAISSLIIKKIIETAKNNYTTLISERLNTVTVENDLLKYIKNKIPVQGRLG
NFKLDTAEFEKKLNTILFVLNESNLAQRFSILVA*KHF*LKERPIDPIYVNILDDNSYSTLEGFNISSQGSN
DFQGQLLESSYFEKIESNALRAFIKICPRNGLLYNAIYRNSKNYLNNIDLEDKKTTSKTNVSYPCSKSLV
PRGSQALLNGCIEVENKDLFLISNKDSLNDINLSEEKIKPETTVFFKDKLPPQDITLSNYDFTEANSIPSI
SQQNILERNEELYEPIRNSLFEIKTIYVDKLTTFHFLEAQNIDESIDSSKIRVELTDSVDEALSNPNKVYS
PFKNMSNTINSIETGITSTYIFYQWLRSIVKDFSDETGKIDVIDKSSDTLAIVPYIGPLLNIGNDIRHGDFV
GAIELAGITALLEYVPEFTIPILVGLEVIGGELAREQVEAIVNNALDKRDQKWAEVYNITKAQWWGTIHL
QINTRLAHTYKALSRQANAIKMNMEFQLANYKGNIDDKAKIKNAISETEILLNKSVEQAMKNTEKFMIKL
SNSYLTKEMIPKVQDNLKNFDLETKKTLDKFIKEKEDILGTNLSSSLRRKVSIRLNKNIAFDINDIPFSEF
DDLINQYKNEIRSGMQVQLVESGGALVQPGGSLRLSCAASGFPVNRYSMRWYRQAPGKEREWVAG
MSSAGDRSSYEDSVKGRFTISRDDARNTVYLQMNSLKPEDTAVYYCNVNVGFEYWGQGTQVTVSS
KSGGYPYDVPDYA

Red: GST, Orange: Thrombin cleavage site, Blue: TTP, Gray: BoNT/X LC (*inactivating mutations), Dark Gray: BoNT/X TD, Green: GFPnb, Purple: HA, Brown: V5

**GST-V5-TTP- BoNT/X-EGFRnb**

MSPILGYWKIKGLVQPTRLLLEYLEEKYEEHLYERDEGDKWRNKKFELGLEFPNLPYYIDGDVKLTQS
MAIIRYIADKHNMLGGCPKERAEISMLEGAVLDIRYGVSRIAYSKDFETLKVDFLSKLPEMLKMFEDRL
CHKTYLNGDHVTHPDFMLYDALDVVLYMDPMCLDAFPKLVCFKKRIEAIPQIDKYLKSSKYIAWPLQG
WQATFGGGDHPPKSDLVPRGSGKPIPNPLLGLDSTKGSGSTSGSGTGPITINNFRYSDPVNNDTIIM
MEPPYCKGLDIYYKAFKITDRIWIVPERYEFGTKPEDFNPPSSLIEGASEYYDPNYLRTDSDKDRFLQT
MVKLFNRIKNNVAGEALLDKIINAIPYLGNSYSLLDKFDTNSNSVSFNLLEQDPSGATTKSAMLTNLIIFG
PGPVLNKNEVRGIVLRVDNKNYFPCRDGFGSIMQMAFCPEYVPTFDNVIENITSLTIGKSKYFQDPALL
LMHELIHVLHGLYGMQVSSHEIIPSKQEIYMQHTYPISAEELFTFGGQDANLISIDIKNDLYEKTLNDYKA
IANKLSQVTSCNDPNIDIDSYKQIYQQKYQFDKDSNGQYIVNEDKFQILYNSIMYGFTEIELGKKFNIKT
RLSYFSMNHDPVKIPNLLDDTIYNDTEGFNIESKDLKSEYKGQNMRVNTNAFRNVDGSGLVSKLIGLC
KKIIPPTNIRENLYNRTAGSSGGSSGMKLEINKFNYNDPIDGINVITMRPPRHSDKINKGKGPFKAFQVI
KNIWIVPERYNFTNNTNDLNIPSEPIMEADAIYNPNYLNTPSEKDEFLQGVIKVLERIKSKPEGEKLLELI
SSSIPLPLVSNGALTLSDNETIAYQENNNIVSNLQANLVIYGPGPDIANNATYGLYSTPISNGEGTLSEV
SFSPFYLKPFDESYGNYRSLVNIVNKFVKREFAPDPASTLMHQ*LVHVTHNLYGISNRNFYYNFDTGKI
ETSRQQNSLIFEELLTFGGIDSKAISSLIIKKIIETAKNNYTTLISERLNTVTVENDLLKYIKNKIPVQGRLG
NFKLDTAEFEKKLNTILFVLNESNLAQRFSILVA*KHF*LKERPIDPIYVNILDDNSYSTLEGFNISSQGSN
DFQGQLLESSYFEKIESNALRAFIKICPRNGLLYNAIYRNSKNYLNNIDLEDKKTTSKTNVSYPCSKSLV
PRGSQALLNGCIEVENKDLFLISNKDSLNDINLSEEKIKPETTVFFKDKLPPQDITLSNYDFTEANSIPSI
SQQNILERNEELYEPIRNSLFEIKTIYVDKLTTFHFLEAQNIDESIDSSKIRVELTDSVDEALSNPNKVYS
PFKNMSNTINSIETGITSTYIFYQWLRSIVKDFSDETGKIDVIDKSSDTLAIVPYIGPLLNIGNDIRHGDFV
GAIELAGITALLEYVPEFTIPILVGLEVIGGELAREQVEAIVNNALDKRDQKWAEVYNITKAQWWGTIHL
QINTRLAHTYKALSRQANAIKMNMEFQLANYKGNIDDKAKIKNAISETEILLNKSVEQAMKNTEKFMIKL
SNSYLTKEMIPKVQDNLKNFDLETKKTLDKFIKEKEDILGTNLSSSLRRKVSIRLNKNIAFDINDIPFSEF
DDLINQYKNEIQVKLEESGGGSVQTGGSLRLTCAASGRTSRSYGMGWFRQAPGKEREFVSGISWRG
DSTGYADSVKGRFTISRDNAKNTVDLQMNSLKPEDTAIYYCAAAAGSAWYGTLYEYDYWGQGTQVT
VSSSGGYPYDVPDYA

Red: GST, Orange: Thrombin cleavage site, Blue: TTP, Gray: BoNT/X LC (*inactivating mutations), Dark Gray: BoNT/X TD, Green: EGFRnb, Purple: HA, Brown: V5

**GST-BTP-BoNT/X-EGFRnb**

MSPILGYWKIKGLVQPTRLLLEYLEEKYEEHLYERDEGDKWRNKKFELGLEFPNLPYYIDGDVKLTQS
MAIIRYIADKHNMLGGCPKERAEISMLEGAVLDIRYGVSRIAYSKDFETLKVDFLSKLPEMLKMFEDRL
CHKTYLNGDHVTHPDFMLYDALDVVLYMDPMCLDAFPKLVCFKKRIEAIPQIDKYLKSSKYIAWPLQG
WQATFGGGDHPPKSDLVPRGSPFVNKQFNYKDPVNGVDIAYIKIPNAGQMQPVKAFKIHNKIWVIPE
RDTFTNPEEGDLNPPPEAKQVPVSYYDSTYLSTDNEKDNYLKGVTKLFERIYSTDLGRMLLTSIVRGIP

FWGGSTIDTELKVIDTNCINVIQPDGSYRSEELNLVIIGPSADIIQFECKSFGHEVLNLTRNGYGSTQYIR
FSPDFTFGFEESLEVDTNPLLGAGKFATDPAVTLAHELIHAGHRLYGIAINPNRVFKVNTNAYYEMSGL
EVSFEELRTFGGHDAKFIDSLQENEFRLYYYNKFKDIASTLNKAKSIVGTTASLQYMKNVFKEKYLLSE
DTSGKFSVDKLKFDKLYKMLTEIYTEDNFVKFFKVLNRKTYLNFDKAVFKINIVPKVNYTIYDGFNLRNT
NLAANFNGQNTEINNMNFTKLKNFTGLFEFYKLLCVRGIITSKTKSLDKGYNKAKGSSGGSSGMKLEI
NKFNYNDPIDGINVITMRPPRHSDKINKGKGPFKAFQVIKNIWIVPERYNFTNNTNDLNIPSEPIMEADAI
YNPNYLNTPSEKDEFLQGVIKVLERIKSKPEGEKLLELISSSIPLPLVSNGALTLSDNETIAYQENNNIVS
NLQANLVIYGPGPDIANNATYGLYSTPISNGEGTLSEVSFSPFYLKPFDESYGNYRSLVNIVNKFVKRE
FAPDPASTLMHQ*LVHVTHNLYGISNRNFYYNFDTGKIETSRQQNSLIFEELLTFGGIDSKAISSLIIKKII
ETAKNNYTTLISERLNTVTVENDLLKYIKNKIPVQGRLGNFKLDTAEFEKKLNTILFVLNESNLAQRFSIL
VA*KHF*LKERPIDPIYVNILDDNSYSTLEGFNISSQGSNDFQGQLLESSYFEKIESNALRAFIKICPRNG
LLYNAIYRNSKNYLNNIDLEDKKTTSKTNVSYPCSKSLVPRGSQALLNGCIEVENKDLFLISNKDSLNDI
NLSEEKIKPETTVFFKDKLPPQDITLSNYDFTEANSIPSISQQNILERNEELYEPIRNSLFEIKTIYVDKLT
TFHFLEAQNIDESIDSSKIRVELTDSVDEALSNPNKVYSPFKNMSNTINSIETGITSTYIFYQWLRSIVKD
FSDETGKIDVIDKSSDTLAIVPYIGPLLNIGNDIRHGDFVGAIELAGITALLEYVPEFTIPILVGLEVIGGEL
AREQVEAIVNNALDKRDQKWAEVYNITKAQWWGTIHLQINTRLAHTYKALSRQANAIKMNMEFQLAN
YKGNIDDKAKIKNAISETEILLNKSVEQAMKNTEKFMIKLSNSYLTKEMIPKVQDNLKNFDLETKKTLDK
FIKEKEDILGTNLSSSLRRKVSIRLNKNIAFDINDIPFSEFDDLINQYKNEIQVKLEESGGGSVQTGGSLR
LTCAASGRTSRSYGMGWFRQAPGKEREFVSGISWRGDSTGYADSVKGRFTISRDNAKNTVDLQMN
SLKPEDTAIYYCAAAAGSAWYGTLYEYDYWGQGTQVTVSSSGGYPYDVPDYA

Red: GST, Orange: Thrombin cleavage site, Blue: BTP, Gray: BoNT/X LC (*inactivating mutations), Dark Gray: BoNT/X TD, Green: EGFRnb, Purple: HA

(3) *An Overview of HEK-293 Cell Line*. https://www.beckman.com/resources/product-applications/lead-optimization/cell-line-development/human-embryonic-kidney-293 (accessed 1/28).
